# Colorectal cancer risk in bowel adenomas based on lifestyle exposures, tissue preconditioning and DNA methylation

**DOI:** 10.1101/2021.08.26.457788

**Authors:** Jaim Sutton, Morgan Moorghen, Lai Mun Wang, Christina Thirlwell, Christodoulos Pipinikas, Attila Lorincz

## Abstract

**Background:** Colorectal cancer (CRC) is associated with patient demographics, lifestyle exposures and molecular alterations. However, it is not possible to determine which adenomas will progress to CRC, as ethically it is unacceptable to leave and follow adenomas. We hypothesised that certain lifestyle exposures at high levels could precondition exposed bowel tissue by changing and aging it, increasing the risks of deleterious DNA methylation and genetic alterations. We used a novel study design comparing adenomas with concurrent CRC (thus more likely exposed to deleterious lifestyle effects) to single adenomas in bowels with no history of CRC; we called these high (HR) and low-risk (LR) adenomas respectively.

**Methods:** We carried out a discovery and replication epigenome-wide association study (EWAS) on 106 HR and 111 LR adenomas, profiled with MethylationEPIC BeadChips. In order, to identify differentially methylated positions (DMP), regions (DMR), and DNAm (DNAmethylation) lifestyle exposures and risks, with adjustment for confounders, and gene ontology (GO) and pathway enrichment. Then, two open-source gene expression omnibus (GEO) validation datasets (52, 57 and 49, 48 HR and LR normal bowel tissues respectively) were analysed for these DNAm lifestyle exposures and risks, with adjustment for confounders.

**Results:** Our EWAS found 5 Bonferroni significant DMPs with absolute delta betas ≥ 5%, and 14 significant DMRs with absolute mean DMR delta betas ≥ 5%, replicated in the *GPX7, RGS3* and *TMEM135* cancer-associated genes. DNAm high alcohol exposures were strongly associated with increased risk of HR adenomas (odds ratio (OR) per standard deviation (SD) = 2.16 (95% confidence interval (CI) 1.55 - 3.09, p-value = 9.7 × 10^-6^)). In the validation datasets, DNAm high alcohol (ORperSD = 2.12 (95% CI 1.35 - 3.55, p-value = 2.0 × 10^-3^) and ORperSD = 1.79 (95% CI 1.14 - 2.96, p-value = 1.7 × 10^-2^)), and high body mass index (BMI) exposures (ORperSD = 1.72 (95% CI 1.13 - 2.73, p-value = 1.5 × 10^-2^)) were associated with increased risk of HR normal bowel tissues.

**Conclusions:** High alcohol and BMI exposures may precondition normal bowel tissues and adenomas for increased risk of DNA methylation alterations associated with CRC progression. The DNAm exposure signatures and our newly identified genes may be useful epigenetic biomarkers for CRC prevention.

## Background

CRC is the third most common cancer in males and females in the United Kingdom (UK) (1,2). It is a multifactorial disease thought to be caused by a combination of patient demographics, molecular alterations and lifestyle factors (3–5). Notably, numerous studies have found that high levels of alcohol consumption, BMI and tobacco smoking can be associated with increased risk of CRC (6–9) and poorer survival (10–12). Prevention is an important method for reducing CRC progression, and currently relies on the identification and removal of early bowel lesions such as adenomas, which are dysplastic growths in the bowel composed of abnormal cells with characteristic nuclear and cytoplasmic aberrations (13). As adenomas can progress into CRC through an adenoma to carcinoma sequence, that is commonly thought to start in abnormal epithelial stem cells, which grows into a small adenoma, then a larger adenoma and then to invasive cancer (14). Traditionally, this adenoma to carcinoma sequence is thought to occur through the selection of specific cancer gene alteration sweeps with each increase in adenoma size, low to high grade dysplasia histology and tubular to villous architecture stage (15,16).

Importantly, not all adenomas will progress into CRC, and it’s difficult to predict those that will versus those that do not. As, it is unethical to leave adenomas in patients’ bowels to follow their natural history and molecular changes, and cell culture and animal model systems are not able to fully define all CRC drivers. Therefore, current clinical, histopathological and somatic molecular predictors of CRC progression have poor accuracy. However, recent studies have started to identify putative cancer driver gene alterations in normal tissues that may clonally persist throughout pre-cancer lesion growth to invasive cancer, and therefore be sensitive biomarkers for cancer progression (17–20). Notably, such cancer gene alterations have been found in normal bowel mucosa epithelial cells (21), and also in skin, lung and gastrointestinal normal tissues that are more exposed to environmental and lifestyle exposures (20,22). Furthermore, a recent study has found that carcinogenic chemical exposures can also cause genetic alterations in mouse normal tissue that may create a high-risk tissue environment for tumour growth and progression (23).

Epigenetic changes are associated with lifestyle exposures and can be detected as altered DNA methylation patterns in blood and tissues. Notably, EWASs have been carried out that have identified DNAm signatures associated with high alcohol consumption, high BMI and current smoking behaviour exposures. These studies encompass genome-wide cytosine-phosphate-guanine (CpG) methylation differences in the extreme levels of each exposure, compared to those without the exposure (24–26). Additionally, high alcohol and current smoking lifestyle exposure effects on DNA methylation have been found in normal and cancer tissues, especially in individuals more exposed to these lifestyle factors (27–29). Furthermore, a recent study has also found DNA methylation alterations in CRC, postulated to have originated in normal bowel tissue, and then maintained during adenoma to CRC progression (30).

We hypothesised that high levels of certain lifestyle exposures could precondition susceptible bowel tissue, by changing and aging these tissues, thereby increasing DNA methylation alterations and associated risk of CRC progression. Then, that in our novel case-control study design, patients’ adenomas in a bowel environment with concurrent cancer are more likely subjected to high levels of deleterious lifestyle exposures compared to single adenomas in a bowel environment with no history of CRC (called HR and LR bowel adenomas, respectively), and may help determine DNA methylation biomarkers for CRC progression. We, therefore, carried out a EWAS in HR compared to LR bowel adenomas in discovery and stratified study-site replication cohorts. The samples were profiled with Illumina MethylationEPIC BeadChips to identify DMPs, DMRs, and DNAm lifestyle exposure levels and association with risk of HR bowel adenomas, with adjustment for confounders, and then GO and pathway enrichment. To compare our results to a partially similar validation study, HR and LR normal bowel tissue GEO DNA methylation datasets were then downloaded and also analysed for DNAm lifestyle exposure levels and association with risk of HR normal bowel tissues (31).

## Methodology

### Study design

The EWAS consisted of 106 HR and 111 LR patients’ bowel adenomas collected retrospectively from formalin-fixed paraffin-embedded (FFPE) tissue blocks at St Mark’s Hospital in London and the John Radcliffe Hospital in Oxford, England, as part of the CRC-adenoma-normal (CAN) tissue and clinical data collection (Figure 1). Only sporadic adenomas graded according to the World Health Organization (WHO) tumour classification guidelines (4,32), were included in this study. The adenomas were then further classified from these guidelines, as either having low-grade dysplasia (LGD) or high-grade dysplasia (HGD) histology and tubular adenoma (TA) or tubulovillous adenoma (TVA) or villous adenoma (VA) architecture (32). Patients were excluded if they were under 18 years of age, had known inherited CRC syndromes, inflammatory bowel disease (IBD), and if they had undergone neoadjuvant therapy. An adenoma was classified as HR, if it came from a patients’ bowel environment that contained at least one adenoma ≥ 8 mm with a fully separate (both lesions bordered by morphologically normal tissue) concurrent CRC, identified from 2010 to the end of 2015 in their hospital pathology records. While an adenoma was classified as LR if it came from a patients’ bowel environment that contained a single adenoma ≥ 8 mm without any other concurrent neoplasia (adenomas, serrated lesions and CRCs), with no CRCs in their history and at least 4 years of follow-up, identified from 2010 to the end of 2012 from their hospital pathology records. It is important to note that if the LR adenomas contained HGD on histology, we retained their classification, but this was adjusted for in future analyses. Additionally, the HR bowel adenomas were originally individually matched to LR bowel adenomas by age at diagnosis, gender, study-site and adenoma lesion-size. However, some HR bowel adenomas were then later excluded, specifically if they came from bowels that also contained concurrent microsatellite instability positive (MSI +) CRCs, as recorded in their hospital pathology records. This was in-case these concurrent MSI + CRCs, which are thought to be associated with inflammation, directly caused a HR bowel environment that influenced the DNA methylation alterations in the adenomas and biased the analysis. Additionally, for each patient, their age, gender, study-site and adenoma bowel-side, lesion-size, dysplasia and architecture characteristics were also collated from their hospital pathology records. Then, these characteristics were compared between the patients with HR and LR bowel adenomas to identify any differences, using the student’s t-test or chi-square test for the continuous and categorical variables respectively. The study was approved by proportional ethical review from the Northern Ireland ethics committee, Integrated Research Application System (IRAS) project identification 152955, approval reference 15/NI/0142.

**Figure 1.**
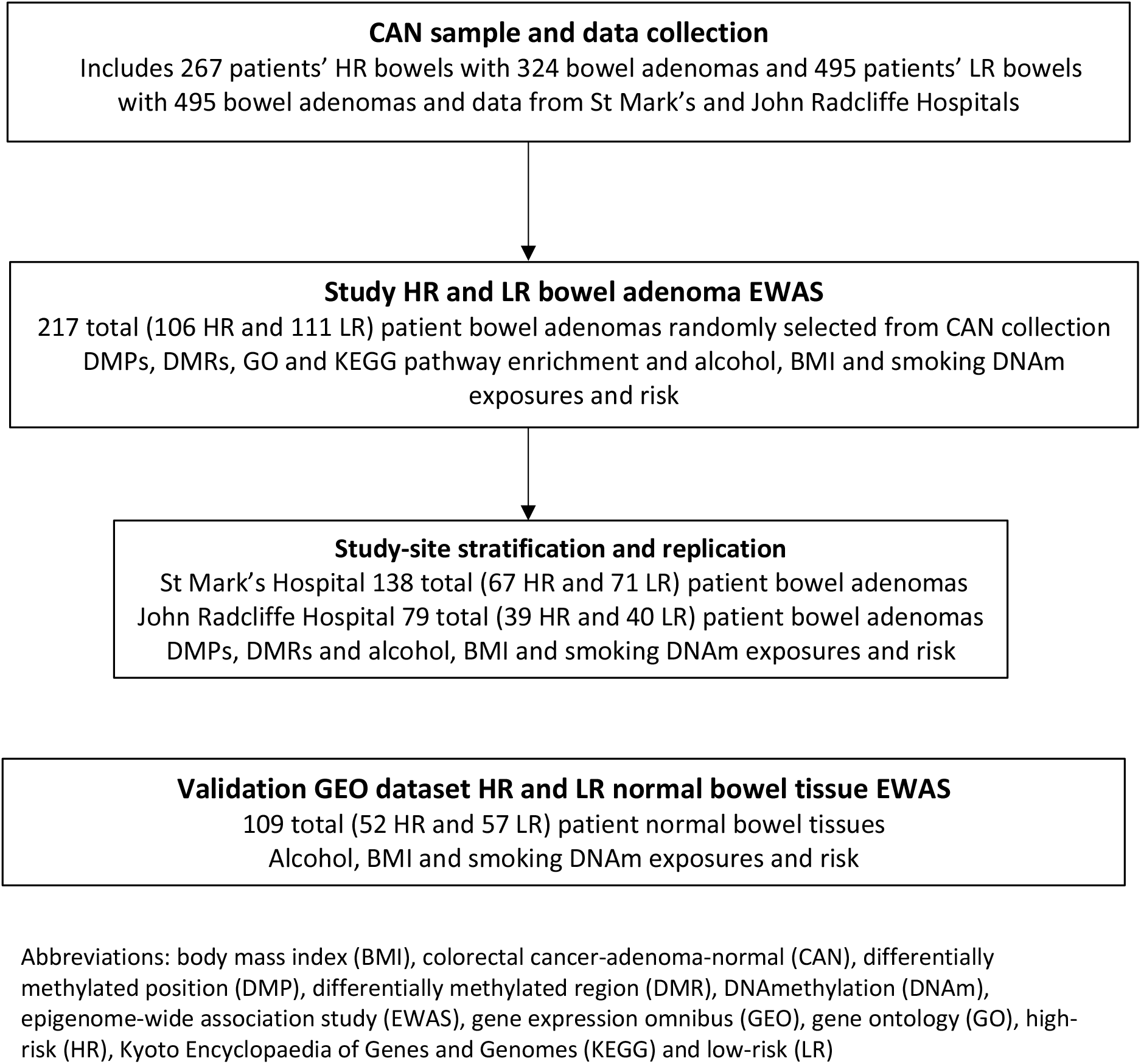
Flow diagram of datasets and analyses. The flowchart outlines the study and validation datasets analysed, and includes sample numbers, and types of analyses.

### DNA methylation profiling and quality control

Adenoma tissue was macro-dissected from pathologist annotated FFPE tissue sections for each patient. Sample DNA was then extracted and bisulfite converted using the Zymo EZ DNA Methylation Kit. Then profiled by the Illumina MethylationEPIC BeadChips at University College London (UCL) Genomics, Institute of Child Health, in London using standard Illumina protocols. The EPIC DNA methylation data was then pre-processed through quality control (QC) steps to produce pre-processed sample beta values, mainly through The Chip Analysis Methylation Pipeline (ChAMP) and minfi pipelines (33–35). These QC steps included single-nucleotide polymorphism (SNP), cross-hybridisation and X and Y chromosome probe filtering (36,37), gender checking, omicsPrint sample relationship genotyping (38), beta-mixture quantile normalization (BMIQ) (39), singular value decomposition (SVD) identification of batch effects through dimension reduction, and ComBat batch effect correction (40,41). From the pre-processed DNA methylation data, the sample beta values were then calculated, by dividing each CpG site M intensity value by their combined target CpG site unmethylated (U) and methylated (M) intensity values (beta value = M/(M + U)). Where, the beta values are continuous variables between 0 (no methylation) and 1 (completely methylated). The sample beta values were then used in the following analyses.

### High and low-risk bowel adenoma epigenome-wide association study

#### Differentially methylated positions

DMP analysis was then carried out using the Confounder Adjusted Testing and Estimation (CATE) package (42), with adjustment for age, gender, study-site and adenoma bowel-side, lesion-size, dysplasia, architecture and 19 latent confounders. The CATE package used a linear model to test for DMPs between the HR and LR bowel adenomas in multiple individual CpG site beta values. It then also identified latent confounders not correlated with case-control status, by surrogate variable analysis (SVA), that were included and adjusted for in the model (42,43). For each model, the DMP p-values were plotted against their expected p-values in quantile-quantile (Q-Q) plots and lambda values calculated, to visualise and identify any DMP p-value genomic inflation. A lambda value of 1.00 to 1.15 represented no or acceptable genomic inflation, respectively. While a lambda value above this, represented high genomic inflation and increased likelihood of DMP false positives, most likely due to un-accounted systematic differences between the HR and LR bowel adenomas. However, any models with high lambda values and inflated DMP p-values were corrected using the BACON (BC) package, that is especially designed to control for EWAS inflation (44). The DMP p-values were then adjusted for multiple testing, as the MethylationEPIC BeadChip includes many individual CpG site tests. With, the conservative Bonferroni and/or less conservative Benjamini-Hochberg false discovery rate (BH FDR) multiple testing correction methods used to determine significant DMP delta beta values with p-values < 7 × 10^-8^ and q-values < 0.05, respectively (45,46). The DMP delta beta values were calculated from the mean CpG site beta value of all HR bowel adenoma samples minus the mean CpG site beta value of all LR bowel adenomas.

#### Differentially methylated regions

DMR analysis was then carried out using the DMRcate package, with adjustment for age, gender, study-site and adenoma bowel-side, lesion-size, dysplasia, architecture and with/without principal components (PC) 1 and 2 (47). DMR analysis identifies regional CpG differences for groupings of CpG sites in the same direction. The DMRcate package computed a kernel-smoothed estimate against a null comparison to identify DMR mean delta values between the HR and LR bowel adenomas, across multiple CpG site beta values with a minimum of 7 CpG site probes. DMRs with Stouffer p-values < 0.05 and smoothed estimate FDR q-values < 0.05, were then used to determine significant DMR mean delta values. A Stouffer value is a summary transformation of individual CpG site FDRs used to rank DMR significance, and PCs 1 and 2 measured the variability in the dataset above 5%. In both the DMP and DMR analyses, positive and negative effect values represent hypermethylation and hypomethylation of the HR compared to the LR bowel adenomas, respectively.

#### Gene ontology and pathway enrichment

GO and Kyoto Encyclopedia of Genes and Genomes (KEGG) pathway enrichment analysis was then carried out using the MissMethyl package, to identify significantly enriched functions and biological pathways for the DMP findings with BH FDR p-values < 0.05 and then for p-values < 0.05. The MissMethyl package was used, as it takes into account the number of CpG site probes per gene in the MethylationEPIC BeadChip during analyses (48).

#### DNAmethylation lifestyle exposures

In, a previously published study, DNAm exposure signatures for overweight/obese BMI, moderate to heavy alcohol consumption and current smoking behaviour were identified from genome-wide CpG methylation differences in the extreme levels of each exposure, compared to those without the exposure. These DNAm high BMI, high alcohol and current smoking exposure signatures consisted of 1109, 450 and 233 CpG sites and corresponding regression coefficients, respectively. Additionally, these DNAm exposure signatures each consisted of a combination of hypermethylated and hypomethylated CpG sites. Notably, the DNAm high alcohol exposure signature included CpG sites for the *SLC7A11* gene, previously found in the literature to be associated with DNAm alterations representing high alcohol consumption (49). We then used these pre-defined CpG site specific DNAm exposure signatures, to further calculate DNAm BMI, alcohol and smoking exposure levels in our patients’ HR and LR bowel adenomas. Where, for each exposure and patient, a DNAm exposure score was calculated from the total sum of each of their DNAm signature CpG site beta values multiplied by their corresponding published DNAm signature regression coefficients. The regression coefficients were used to weight their corresponding CpG site beta values in our study according to their importance in each pre-defined DNAm exposure signature. DNAm exposure scores were then standardized to z-scores and used in unconditional logistic regression analyses, to determine if the DNAm lifestyle exposure levels measured in our patients’ adenoma bowel tissue were associated with increased risk of HR bowel adenomas. In these predictive models, patients’ HR and LR bowel adenoma status was used as the outcome variable and their EWAS measured adenoma DNAm exposure z-scores were used as the predictor variables with adjustment for age, gender, study-site and adenoma bowel-side, lesion-size, dysplasia, architecture and with/without PCs 1 and 2. These models were classified as significant if they had p-values < 0.05, as multiple testing correction was not required for these pre-defined DNAm signatures, as they are classified as a single test.

#### Stratification and study-site replication

Our 106 HR and 111 LR bowel adenoma EWAS dataset, was then also then stratified into St Mark’s Hospital (total 138 bowel adenomas; 67 HR and 71 LR) and John Radcliffe Hospital (total 79 bowel adenomas; 39 HR and 40 LR) study-sites (Figure 1). Then, for each study-site dataset the following analyses were carried out. Firstly, DMP analysis using the CATE package (42), with adjustment for age, gender, and adenoma bowel-side, lesion-size, dysplasia, architecture and 8 latent confounders. Then, DMR analysis using the DMRcate package, with adjustment for age, gender, and adenoma bowel-side, lesion-size, dysplasia, architecture and with/without PCs 1 and 2 (47).

The previously calculated HR and LR bowel adenoma DNAm lifestyle exposure z-scores, where then used in study-site unconditional logistic regression analyses, with adjustment for age, gender and adenoma bowel-side, lesion-size, dysplasia, architecture and with/without PCs 1 and 2. For all the study-site analyses, the same correction methods and p-value significant thresholds were used as in the combined analyses. Apart from the John Radcliffe Hospital study-site model analyses, where adenoma bowel-side was not included and adjusted for, due to an imbalance between the cases and controls.

### Validation in high and low-risk normal bowel tissue

The open-source Gene Expression Omnibus (GEO) HR and LR normal bowel tissue datasets (GSE132804 - GPL21145 and GPL13534) (31), were downloaded from www.ncbi.nih.gov/gds and used for validation. This dataset included similarly classified HR and LR normal bowel tissue patient samples to the HR and LR bowel adenomas. Where, HR was defined as normal bowel tissue in a patients’ bowel with concurrent CRC, and LR as normal bowel tissue in a patients’ bowel with no concurrent adenomas; there was no data on adenomas in this GEO dataset. From these datasets, Illumina MethylationEPIC BeadChip and clinical data from 109 normal bowel tissue samples (52 HR and 57 LR) and Illumina Methylation450K BeadChip and clinical data from 97 normal bowel tissue samples (49 HR and 48 LR), collected from 2017 to 2019 and 2012 to 2016 respectively, were selected. We, then applied the same pre-process (probe filtering, normalisation and relevant batch effects correction) and DNAm exposure and risk association analyses steps to these HR and LR normal bowel tissue datasets, as previously described for our HR and LR bowel adenoma study, so they could be more comparable. Notably, for the DNAm exposure analyses, we used the previously described published study (50) that identified the pre-defined DNAm high BMI, high alcohol and current smoking exposure signatures, to further calculate these DNAm exposure levels in patients’ normal bowel tissue (Figure 1). Where, for each exposure and patient, a DNAm exposure score was calculated from the total sum of each of their DNAm signature CpG site beta values multiplied by their corresponding published DNAm signature regression coefficients. These DNAm lifestyle exposure scores were then standardized to z-scores for subsequent analyses and exposure comparison. Unconditional logistic regression analyses were then carried out to determine if the DNAm lifestyle exposure levels measured in patients’ normal bowel tissue were associated with risk of HR normal bowel tissue. In these models, patients’ HR and LR normal bowel tissue status was used as the outcome variable and their normal bowel tissue DNAm exposure z-scores as the predictor variable with adjustment for age, gender and with/without PCs 1 and 2. These models were classified as significant if they had p-values < 0.05. Additionally, both HR and LR normal bowel tissue datasets only included left normal bowel-sided tissue. However, these should compare well with the HR and LR bowel adenoma dataset, which was mostly left-bowel-sided.

### Statistical analyses

A flowchart outlining all of the study and GEO datasets and analyses are shown in Figure 1, and further dataset details can be found in the supplementary section and in the previously published paper. All statistical analyses were carried out with R version 3.6 and annotation with the IlluminaHumanMethylationEPICanno.ilm10b2.h19 package and GRCh37/hg19 build.

## Results

### Study population characteristics

The HR and LR bowel adenoma study characteristics are described in Table 1. Our study included 64% (n = 138) and 36% (n = 79) patients from the St Mark’s and John Radcliffe Hospitals, respectively. The number of left versus right-sided bowel specimens was significantly different in the HR compared to LR bowel adenomas (chi-square test p-value < 0.001). With overall more adenomas detected on the left bowel-side but with a relative enrichment for HR bowel adenomas on the right bowel-side. Further details of the patients’ HR bowels are also described in Table S1. Notably, it was found that 62% of the HR bowels contained 1 to 3 adenomas concurrently.

**Table 1.**
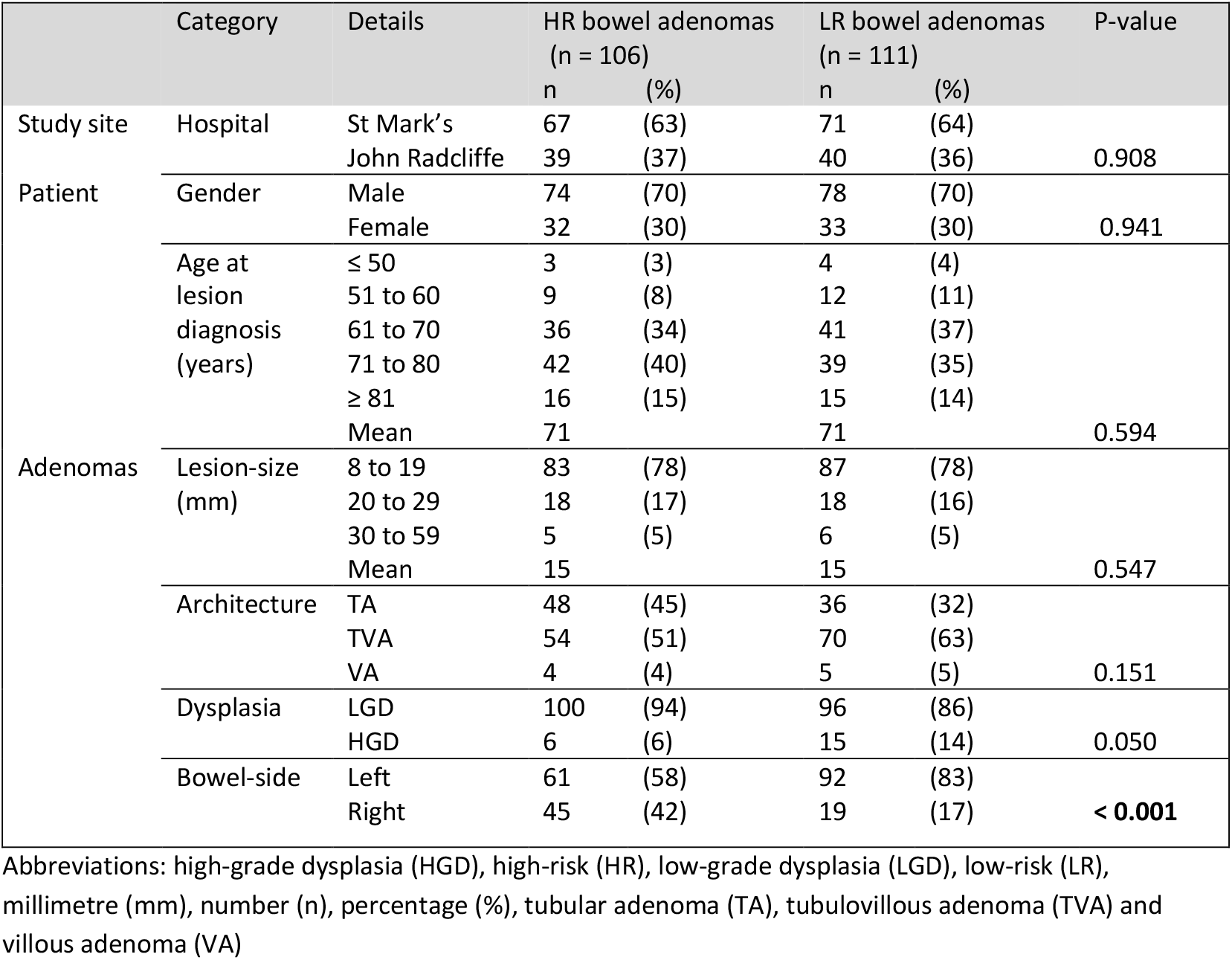
High and low-risk bowel adenoma epigenome-wide association study characteristics. Clinical, histopathological and demographic characteristics for the patients and high-risk (HR) and low-risk (LR) bowel adenomas are shown. The student’s t-test and chi-square test was used to test for differences between continuous data and categorical data, respectively. Significant p-values < 0.05 are highlighted in bold.

### Methylation BeadChip quality control

The methylation levels of 866,836 CpG positions across the genome were measured using the MethylationEPIC BeadChip in the HR and LR bowel adenomas, then pre-processed through QC steps. During these steps CpG positions were removed that failed pre-specified QC criteria. The final number of CpG positions per sample were 713,255 with the following probes removed: 39,117 that failed hybridisation at detection p-value > 0.01, 5,537 with a bead count < 3 in at least 5% of samples, 2,457 non-CpG probes, 91,005 with SNPs close to the 3’ end, 11 that could bind to multiple regions in the genome, and 14,566 in the X and Y chromosomes (34,35). Sample well position, BeadChip and plate number were found to be significant batch effects, and the sample beta values were then adjusted for these batch effects (40,41). All samples were then found to have good beta density distributions, some case-control clustering, identical clinical and bioinformatic calculated genders and were not related to each other by SNP relationship analysis (38) (Figure S1).

### High and low-risk bowel adenoma epigenome-wide association study analysis

#### Differentially methylated positions

DMP analysis after confounder adjustment and genomic inflation correction, identified 5 DMPs with Bonferroni p-values < 7 × 10^-8^ and absolute delta beta values ≥ 5%, and 95 DMPs with BH FDR q-values < 0.05 and absolute delta beta values ≥ 10% (Figure S3). Genomic inflation correction was able to reduce the model lambda from 2.10 to 1.09, resulting in low levels of DMP genomic inflation and thus relatively decreased false positives (Figure S2). Notably, three of the DMPs that passed Bonferroni significance were found in the previously described *DIAPH3, RGS3* and *TMEM135* cancer-associated genes (51–53). Additionally, the most significant annotated DMP was hypomethylation in the *DIAPH3* gene, which plays a role in actin remodelling, epithelial-mesenchymal transition (EMT) and cancer signalling (delta beta = - 6.7%, BH FDR q-value = 1.6 × 10^-3^). While the annotated DMP with the largest absolute delta beta value was hypomethylation in the *ANK2* gene, which plays a role in the structural part of cytoskeleton and enzyme binding (delta beta = - 8.1%, BH FDR q-value = 2.7 × 10^-2^). Interestingly, most of our highly significant DMP delta beta values were hypomethylated in the HR versus the LR bowel adenomas, and generally spanned widely across the human genome (Figure S3). The DMPs with delta beta values > 5% and BH FDR q-values < 0.05 are shown in Table 3. Notably, some of these tabled DMPs were found to be associated with lifestyle factors, epithelium, adenomas, progression and CRC, and other cancers in the literature. With gene functions encompassing telomere length, EMT, and cancer growth, invasion and signalling pathways (Table S2).

#### Differentially methylated regions

DMR analysis after confounder adjustment, identified 2,074 DMRs, and of these 14 DMRs had absolute mean delta beta values ≥ 5%. The most significant DMR was hypermethylation in the *GPX7* gene, that plays a role in DNA damage and double-strand break protection (mean delta beta = 6.2%, Stouffer p-value = 4.26 × 10^-12^). Additionally, the DMR with the largest absolute mean delta beta value was hypermethylation in the *MCIDAS* gene that plays a role in transcription coactivator activity (mean delta beta = 10.4%, Stouffer p-value = 1.83 × 10^-9^). The DMRs with mean delta beta values > 5% and Stouffer p-values < 0.05, with/without PCs adjustment were similar and mostly hypermethylated for the HR compared to LR bowel adenomas (Tables S3 and 4). Notably, some of these tabled DMRs were found to be associated with lifestyle factors, epithelium, progression and CRC, and other cancers in the literature. With gene functions encompassing transcriptional factor activity, DNA damage, immunity, and cancer signalling pathways (Table S4).

#### Gene ontology and pathway enrichment

GO and KEGG pathway enrichment analysis of the DMPs with significant BH FDR p-values < 0.05, did not find any terms and pathways enriched in the HR bowel adenomas at BH FDR q-values < 0.05, most likely due to low sample size power. However, the GO and KEGG pathway enrichment analysis of the DMPs with significant p-values < 0.05, found many terms and pathways enriched in the HR bowel adenomas at BH FDR q-values < 0.05 (Table S5). Such as terms associated with metabolic, cellular and molecular processes, bacterial invasion of epithelium, metabolism, and cancer and associated signalling. All of which, if aberrantly methylated, could potentially increase the risk of adenoma to CRC progression in HR bowel environments.

### DNA methylation lifestyle exposures in bowel tissue

#### High and low-risk bowel adenoma tissue

In the HR and LR bowel adenoma dataset, the number of CpG sites and beta-values that were available in our data corresponding to the published DNAm exposure signature CpG sites were as follows: 1021/1109, 413/450 and 211/233 for the BMI, alcohol and smoking exposures respectively. DNAm exposure logistic regression analyses, after adjustment for confounders, identified significant associations for DNAm high alcohol and high BMI exposures with the risk of HR bowel adenomas, at p-values < 0.05. Notably, a one SD unit increase in DNAm high alcohol exposure score was found to be strongly significantly associated with an increased risk of HR bowel adenomas (ORperSD = 2.16 (95% CI 1.51 - 3.09, p-value = 9.7 × 10^-6^)). While a one SD unit increase in DNAm high BMI and then current smoking exposure scores were found to be weakly and not significantly associated with a decreased risk of HR bowel adenomas, respectively (Table 2).

**Table 2.** High and low-risk bowel tissue and DNAmethylation lifestyle exposures. DNAmethylation (DNAm) alcohol, body mass index (BMI) and smoking exposure associated odds ratios (ORs) for the risk of high-risk (HR) bowel adenomas, stratified study-site HR bowel adenomas and external HR normal bowel tissue, adjusted for confounders and without/with principal components 1 and 2 are shown. The normal bowel tissue is left bowel-sided and obtained from the gene expression omnibus (GEO) public database. Significant associations are highlighted in bold.

| Datasets | Total (n HR, n LR) | DNAm exposures | ORperSD | 95% CI | P-value |
| --- | --- | --- | --- | --- | --- |
| HR and LR bowel adenomas EWAS datasets |  |  |  |  |  |
| All<br>(EPIC 850K) | 217 (106, 111) | Alcohol | **2.16** | **1.55 - 3.09** | **9.7e-06** |
|  |  |  | **2.12**<sup>1</sup> | **1.51 - 3.04** | **2.7e-05** |
|  |  | BMI | **0.70** | **0.51 - 0.95** | **2.4e-02** |
|  |  |  | **0.72**<sup>1</sup> | **0.52 - 0.99** | **4.6e-02** |
|  |  | Smoking | 0.86 | 0.64 - 1.15 | 3.2e-01 |
|  |  |  | 0.88<sup>1</sup> | 0.65 - 1.19 | 4.2e-01 |
| SM | 138 (67, 71) | Alcohol | **1.85** | **1.24 - 2.84** | **3.5e-03** |
|  |  |  | **1.85**<sup>1</sup> | **1.21 - 2.89** | **5.3e-03** |
|  |  | BMI | 0.84 | 0.57 - 1.24 | 3.9e-01 |
|  |  |  | 0.87<sup>1</sup> | 0.58 - 1.30 | 4.9e-01 |
|  |  | Smoking | 0.89 | 0.61 - 1.28 | 5.3e-01 |
|  |  |  | 0.92<sup>1</sup> | 0.63 - 1.35 | 6.8e-01 |
| JR | 79 (39, 40) | Alcohol | **3.44** | **1.91 - 7.00** | **1.6e-04** |
|  |  |  | **5.24**<sup>1</sup> | **2.57 - 13.01** | **5.1e-05** |
|  |  | BMI | **0.36** | **0.17 - 0.66** | **3.0e-03** |
|  |  |  | **0.37**<sup>1</sup> | **0.17 - 0.70** | **5.4e-03** |
|  |  | Smoking | 0.78 | 0.47 - 1.26 | 9.0e-01 |
|  |  |  | 0.78<sup>1</sup> | 0.46 - 1.29 | 3.4e-01 |
| GEO HR and LR normal bowel tissue EWAS dataset |  |  |  |  |  |
| GLP22145<br>(EPIC 850K) | 109 (52, 57) | Alcohol | **2.12** | **1.35 - 3.55** | **2.0e-03** |
|  |  |  | 1.37<sup>1</sup> | 0.78 - 2.50 | 2.9e-01 |
|  |  | BMI | **1.72** | **1.13 - 2.73** | **1.5e-02** |
|  |  |  | **2.73**<sup>1</sup> | **1.52 - 5.39** | **1.7e-03** |
|  |  | Smoking | **0.59** | **0.38 - 0.90** | **1.7e-02** |
|  |  |  | 0.79<sup>1</sup> | 0.45 - 1.37 | 4.0e-01 |
| GPL13534<br>(450K) | 97 (49, 48) | Alcohol | **1.79** | **1.14 - 2.96** | **1.7e-02** |
|  |  |  | 0.80<sup>1</sup> | 0.39 - 1.61 | 5.4e-01 |
|  |  | BMI | 1.11 | 0.85 - 1.44 | 4.5e-01 |
|  |  |  | 0.70<sup>1</sup> | 0.46 - 1.02 | 7.7e-02 |
|  |  | Smoking | **0.28** | **0.14 - 0.50** | **8.1e-05** |
|  |  |  | 0.49<sup>1</sup> | 0.20 - 1.08 | 9.2e-02 |

Abbreviations: body mass index (BMI), confidence interval (CI), DNAmethylation (DNAm), epigenome-wide association study (EWAS), exponential notation (e), gene expression omnibus (GEO), high-risk (HR), John Radcliffe Hospital (JR), low-risk (LR), number (n), odds ratio (OR), standard deviation (SD) and St Mark's Hospital (SM)
<sup>1</sup> Principal components 1 and 2 adjustment

### Study-site stratification and replication

DMP analyses after confounder adjustment and genomic inflation correction, found significantly replicated DMPs with delta beta values in the same direction in both study-sites, at BH FDR p-values < 0.05 for the larger St Marks Hospital and at p-values < 0.05 for the smaller John Radcliffe Hospital study-sites, and previously in the combined analysis. These replicated DMPs were found in the previous RGS3 and TMEM135 cancer-associated genes (52,53) in a novel *SRP14-AS1* gene, and in the open sea cg22240312 and cg17332384 sites (Tables 3 and S2). Genomic inflation correction was also able to reduce the model lambdas from 1.89 to 1.09 for the St Mark’s Hospital, and 1.62 to 1.15 for John Radcliffe Hospital study-sites (Figure 1). Resulting in low levels of DMP p-value genomic inflation and thus relatively decreased false positives for both study-sites. DMR analyses after confounder adjustment, then found the *GPX7, XKR6, LBX2* genes to be significantly replicated DMRs in both hospital study-sites, with mean delta beta values in the same direction and Stouffer p-values < 0.05, consistent with the earlier combined analysis (Tables 4 and S4).

**Table 3.**
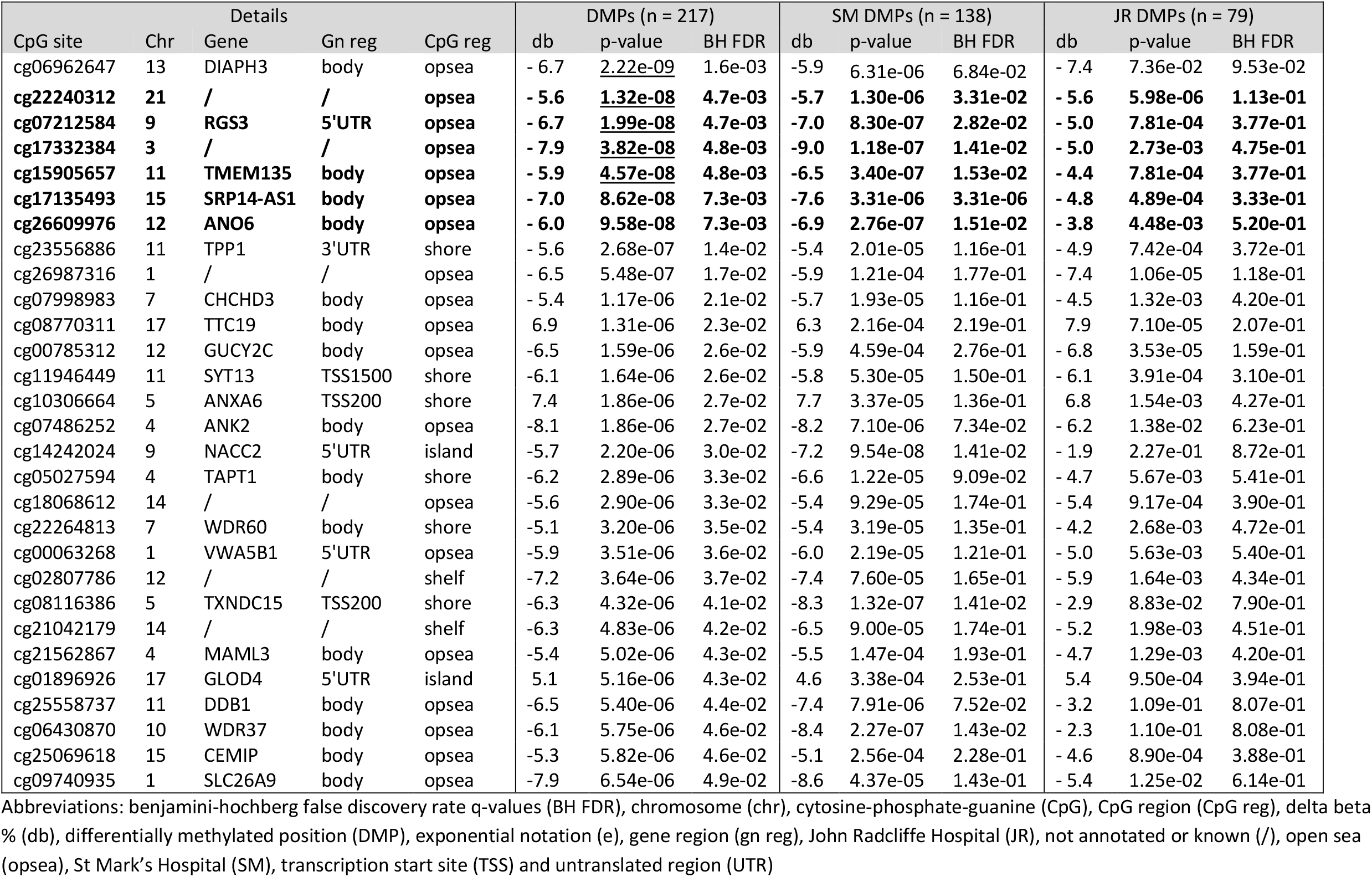
Differentially methylated positions. High-risk (HR) compared to low-risk (LR) bowel adenoma differentially methylated positions (DMPs) adjusted for confounders and genomic inflation corrected, with delta betas values ≥ 5% and Benjamini-Hochberg false discovery rate (BH FDR) q-values < 0.05, are shown. These were ordered by decreasing p-value significance, and then matched to their corresponding stratified St Mark’s Hospital and John Radcliffe Hospital study-site DMP values. The Bonferroni significant p-values < 7 × 10^-8^ after genomic control are underlined and significant replicated study-site DMPs are highlighted in bold.

**Table 4.**
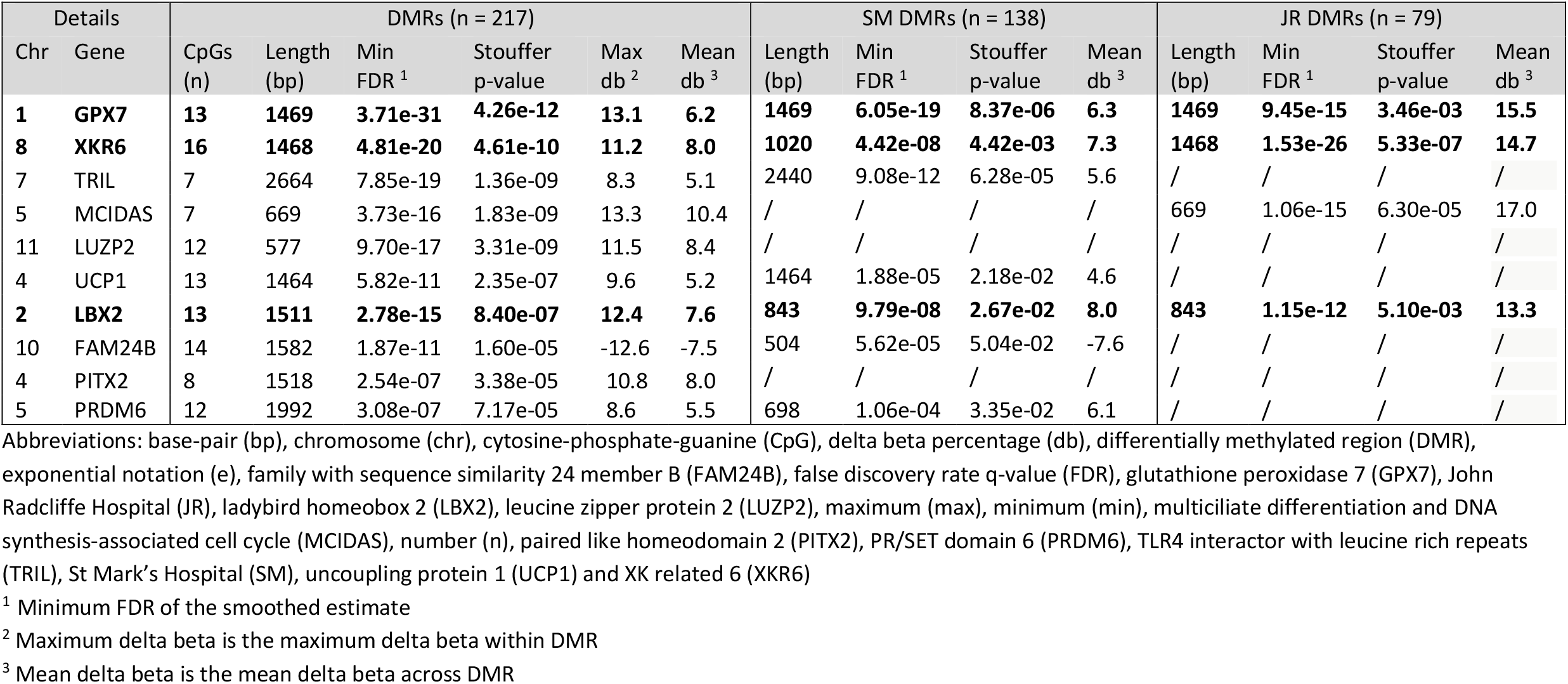
Differentially methylated regions without principal component adjustment. High-risk (HR) compared to low-risk (LR) bowel adenoma differentially methylated regions (DMRs) adjusted for confounders, with mean delta beta values ≥ 5%, Stouffer p-values < 0.05 and minimum of 7 cytosine-phosphate-guanines (CpGs) are shown. These were ordered by decreasing Stouffer p-values and then matched to their corresponding stratified St Mark’s Hospital and John Radcliffe Hospital study-site DMR values. Significant replicated study-site DMRs are highlighted in bold.

Notably strong effects were seen for the DMR in the *GPX7* gene, for both St Marks Hospital (DMR mean delta beta = 6.3%, Stouffer p-value = 8.37 × 10^-6^) and John Radcliffe Hospital study-sites (DMR mean delta beta = 15.5%, p-value = 3.46 × 10^-3^) (Table 4). This was also the only DMR gene to be similarly and significantly replicated in both study-sites, after further PC adjustment (Table S3). Then, the DNAm exposure logistic regression analyses after adjustment for confounders only found DNAm high alcohol exposure to be significantly associated with the risk of HR bowel adenomas at p-values < 0.05, in both study sites and the previous combined analysis. With, a one SD unit increase in DNAm high alcohol exposure score found to be significantly associated with an increased risk of HR bowel adenomas for the St Mark’s Hospital (ORperSD = 1.85 (95% CI 1.24 - 2.84, p-value = 3.5 × 10^-3^)), and John Radcliffe Hospital study-sites (ORperSD = 3.44 (95% CI 1.91 - 7.00, p-value = 1.6 × 10^-4^)). While a one SD unit increase in DNAm high BMI exposure score was found to be associated with a decreased risk of HR bowel adenomas for each study-site, but only significantly for the smaller John Radcliffe Hospital study-site. Then a one SD unit increase in DNAm current smoking exposure score was found to be associated with a decreased risk of HR bowel adenomas for each hospital study-site, but not significantly (Table 2). The study-site DNAm exposure logistic regression analyses with PCs adjustment are also shown in Table 2. Where only the DNAm high alcohol exposure score was found to be significantly associated with an increased risk of HR bowel adenomas, with and without PC adjustment, in both study-sites and the previous combined analysis.

### Validation in high and low-risk normal bowel tissue

In the GEO GSE132804 Illumina MethylationEPIC BeadChip HR and LR normal bowel tissue GPL21145 and GPL13534 validation datasets, the number of CpG sites and beta-values that were available out of the published DNAm exposure signature CpG sites were as follows: 1056/1109, 428/450 and 215/233 for GPL21145, and then 1092/1109, 445/450 and 228/233 for GPL13534, for the BMI, alcohol and smoking exposures respectively. Then, the DNAm exposure logistic regression analyses, after adjustment for confounders, only identified significantly replicated associations for DNAm high alcohol exposure with the risk of normal bowel tissue, at p-values < 0.05, in both validation datasets and all previous study-site analyses. With a one SD unit increase in DNAm high alcohol exposure score found to be significantly associated with an increased risk of HR normal bowel tissues for the GPL21145 (ORperSD = 2.12 (95% CI 1.35 - 3.55, p-value = 2.0 × 10^-^ 3)), and GPL13534 datasets (ORperSD = 1.79 (95% CI 1.14 - 2.96, p-value = 1.7 × 10^-2^)), and in the previous combined and study-site replicated analyses. While a one SD unit increase in DNAm current smoking exposure score was found to be significantly associated with a decreased risk of HR normal bowel tissue in both validation datasets, but only strongly for the GPL13534 dataset (ORperSD = 0.28 (95% CI 0.14 - 0.50, p-value = 8.1 × 10^-5^)), and not significantly in any of the previous combined and study-site replicated analyses. Then, a one SD unit increase in DNAm high BMI exposure score was associated with an increased risk of HR normal bowel tissue in both validation datasets, but only significantly for the GPL21145 dataset (ORperSD = 1.72 (95% CI 1.13 - 2.73, p-value = 1.5 × 10^-2^)) (Table 2). The validation DNAm exposure logistic regression analyses with PCs adjustment for both datasets are also shown in Table 2.

## Discussion

In summary, our HR versus LR bowel adenoma EWAS, found highly significant and replicated DMPs with absolute delta beta values ≥ 5%. We also identified a significantly replicated DMR with absolute mean DMR delta beta values ≥ 5%. These are large and significant effect sizes in what might be diagnostically important and druggable CRC genes. Our results are novel for potentially predicting HR adenomas using molecular methods. Some of our observed effects were seen in genes previously associated with cancer (52–54). Another important result, is that a previously derived and published DNAm high alcohol exposure signature (25), found that DNAm high alcohol bowel exposure levels were significantly associated with increased risk of HR bowel adenomas in our data. While, in the validation dataset, both DNAm high alcohol and high BMI bowel exposure levels were found to be significantly associated with increased risk of HR normal bowel tissue.

It could be, that high levels of DNAm measured lifestyle exposures in a patients’ bowel could precondition the exposed normal and later adenoma tissue by changing and aging this tissue. This could then increase the risk and occurrence of early clonal or later sub-clonal DNA methylation alterations in the epithelium stem cells, other normal and adenoma bowel tissue, creating a high-risk tissue environment for tumour growth and progression. Notably, for high BMI and alcohol tissue exposures, there is evidence for such a proposed cancer progression model in the literature. As, high BMI exposure has been found to be associated with increased risk for breast and colorectal cancers and/or poorer survival (55,56), which originate in tissues more likely to be exposed to obesity-associated metabolites. Additionally, a recent study has identified a transcriptomic metabolic subtype in adjacent normal tissue to breast cancer that was also associated with poor breast cancer prognosis (57). Consequently, in our study the DNAm high BMI scores may be measuring the impact of high levels of obesity-associated metabolites related to changes in metabolic reprogramming, lipid levels and other processes, that could precondition the exposed normal bowel tissue and alter epigenetically controlled differentiation pathways (57,58). High alcohol exposure has also been found to be associated with increased risk and/or poorer survival of head and neck, pancreatic, esophageal and colorectal cancers, that originate in tissues more likely to be exposed to high levels of alcohol metabolites (7,10,11,59,60). Additionally, somatic genetic alcohol mutational signatures have also been found to be associated with cancers likely to be exposed to high levels of alcohol metabolites, and to occur clonally in normal tissue before carcinogenesis (22,61,62). Therefore, high BMI and alcohol exposures could precondition bowel tissue for increased risk of CRC progression and reveal relevant DNA methylation biomarkers.

In our study, the significant differentially methylated positions and regions found between the HR and LR adenomas could be drivers of CRC progression. As many of our observed biomarkers are in plausible effector genes and previously associated with early bowel tissues, cancer progression and risk, in the literature. As for the DMPs, notably the *GUCY2C* gene, a hypermethylated study DMP, was previously found to be associated with obesity and gene signalling silencing in normal bowel epithelial cells, where clonal cancer driver alterations are expected to occur (63,64). Then, the *ANXA6* gene, a hypomethylated study DMP, was postulated to be a HR adenoma biomarker, in a study where patients’ adenomas were classified as HR if had cancer associated copy-number alterations, and then found to express this protein in their stools with high sensitivity (65). While the *TPP1* gene, a hypermethylated study DMP, was previously found to be DNA altered and associated with telomere length dysfunction and increased CRC risk (66). Additionally, the *TMEM135* gene, a hypermethylated study DMP, was previously found to be near a SNP associated with increased breast cancer risk (53). Furthermore, the study-site replicated hypermethylated DMPs in the *DIAPH3* and *RGS3* genes, were previously found to be associated with EMT, which is important in cancer transformation, and with adenoma and cancer signalling pathways, such as the wingless-type (Wnt), and mitogen-activated protein kinase (MAPK) pathways, respectively (51,52).

Then, for the significant DMRs, notably, the *GPX7* gene, a study-site replicated hypermethylation DMR, was previously found to be associated with DNA damage repair and alterations in epithelial cells in normal bowel tissue compared to gastric cancer, again where clonal cancer driver alterations are expected to occur. While, the *MCIDAS* gene, a hypermethylated study DMR with a large effect size, was previously found to be hypermethylated in serrated CRC versus normal bowel tissue (67). Additionally, the *CCL28* gene, a hypermethylated study DMR, was previously found to be associated with immune response and transforming growth factor beta (TCFß) signalling pathway, and then with reduced expression in left CRC versus normal bowel tissue (68). Furthermore, the *PITX2* gene, a hypermethylated study DMR, was previously found to be associated with promoter hypermethylation in CRC versus normal bowel tissue and then reduced expression was also associated with poorer survival (69,70). These positional and regional differentially DNA methylated results were often methylated in a direction that matched their cancer associated literature, and taken in combination, strengthens the evidence that our observed alterations could be due to normal and/or adenoma bowel tissue CRC preconditioning due to high lifestyle bowel exposures, and causally associated with CRC progression. Therefore, these DNA methylation gene alterations could be possible HR adenoma CRC progression biomarkers and target genes suitable for interventions via lifestyle modification and preventative drug interventions.

There are several strengths to our study. Notably, EWAS differential methylation findings are commonly influenced by confounders, which are variables that if distributed unequally between the cases and controls, can bias the findings. Therefore, in this study model analyses were adjusted for patient demographic, adenoma and bowel confounders, by model inclusion of these collected variables. While other relevant latent confounders or PCs were then also included in the models to adjust for unaccounted confounders such as ethnicity, adenoma cell type composition, diagnosis route, and general FFPE block age. Selected findings were also corrected for genomic inflation, adjusted for multiple testing and stratified by study site, to reduce the occurrence of false positives and further increase validity. Additionally, the Infinium Methylation EPIC BeadChip technology has a relatively low error rate and has been shown to compare well with whole-genome bisulfite sequencing (WGBS), therefore our study findings do not need to be cross-validated by another molecular technique (71).

There are some limitations to our study. Firstly, the EWAS study numbers were relatively small and the EPIC methylation BeadChip has low genome coverage, including less than 3% of all CpG sites (72), which is just a fraction of the full methylome. The low coverage would impact the numbers of DMPs and DMRs we discovered but not the validity of their discovered effects. As, there are probably many other relevant DMPs and DMRs related to adenoma to CRC progression that remain to be discovered. Then, model analyses that include PCs, can lead to over adjustment and reduced measurement of the biological effect, especially with small sample sizes. However, in this study, model analyses with and without PCs were both run and presented, with more focus on the latter. Additionally, in the stratified study-site analyses, bowel-side could not be adjusted for the John Radcliffe Hospital dataset. However, only findings that had been originally adjusted for bowel-side in the unstratified analyses were investigated for study-site replication. Furthermore, with this study design, causality can’t be concluded, and therefore the concurrent cancer may be responsible for the HR adenoma DNA methylation alterations. However, this is unlikely, as the study adenomas were checked thoroughly for cancer by the study pathologists, and any patients’ HR bowels with concurrent MSI+ CRC recorded in their hospital pathology records, were excluded, in-case they created a HR bowel environment. It is also expected that any concurrent cancers not previously selected and recorded for MSI testing and status, would have a low or negligible proportion of MSI + typing.

Next, the DNAm lifestyle exposure signatures used in this study were originally identified in blood and may not translate well and be maintained in bowel tissue. However, one published study has found that tissue from alveoli has a similar epigenetic profile with blood-derived samples at CpG sites associated with smoking (28). Early bowel progression tissue would also be expected to be less molecularly altered than in later invasive CRC tissue. Additionally, it is likely that there may be unmeasured environmental and lifestyle exposures in the HR bowel environment that may have also contributed to causing the DNA methylation alterations, some of which could be measured in the future, with further developed lifestyle exposure signatures and assays. Furthermore, the pre-defined DNAm current smoking exposure signature used in this study, may have also shown conflicting results because it is not suited to bowel tissue and may need re-development. Also, due to restrictions in the sample collection where follow-up data was required for the LR bowel adenomas, there was an in-balance of FFPE block age between the HR (2010 to 2015 block age years) and LR bowel adenomas (2010 to 2013 block age years), that made precise FFPE block age adjustment difficult. However, it has been found that there was no significant deterioration of methylation concordance between sample duplicates in archived FFPE tissue blocks up to 10 years of age in one study (73) and generally in other methylation BeadChip array studies (71,74,75). Lastly, additional external validations were not able to be carried out, but would be desirable, when such data is available.

## Conclusions

High levels of alcohol and BMI bowel exposures may precondition normal bowel tissue, and also adenoma bowel tissue, for increased risk of DNA methylation alterations that aid high-risk adenoma growth and CRC progression. The effect of BMI is less clear in our study, but high BMI is probably quite important and has been suggested as a risk factor for CRC in many earlier studies. Our inability to show an obvious association between HR adenomas and high BMI bowel exposure may be more a function of our study power than a lack of actual effects, and/or a stronger effect earlier in CRC progression. Finally, tobacco use did not manifest as an exposure that could be associated with DNA methylation and increased risk of HR bowel adenomas in our study. Overall, our findings could potentially be used in the future to risk stratify normal bowel tissue and adenomas that are more likely to progress to CRC and may be useful biomarkers to aid CRC early detection, screening and prevention. It also brings to light a new study design that may potentially be useful in identifying further HR adenoma molecular biomarkers for CRC progression. However, future investigations are needed to determine if DNAm high BMI and high alcohol exposure bowel tissue levels and the HR adenomas DNA methylation alterations can be further associated with risk of CRC progression.

## Acknowledgments

The authors would like to thank the study participants and study/healthcare staff who have contributed to the study cohorts. As well as, members of the Molecular Epidemiology and Evolution Cancer Laboratories, Barts and the London School of Medicine and Dentistry, Queen Mary and the Medical Genomics Group, Cancer Institute, University College London, University of London for their advice and support during the study. The study dataset was obtained from our larger CRC-adenoma-normal (CAN) tissue and clinical data collection, and collaborations are welcome.

