## Supplementary material for "Colorectal cancer risk in bowel adenomas based on lifestyle exposures, tissue preconditioning and DNA methylation"

### Supplementary Figures and Tables

#### Figure S1

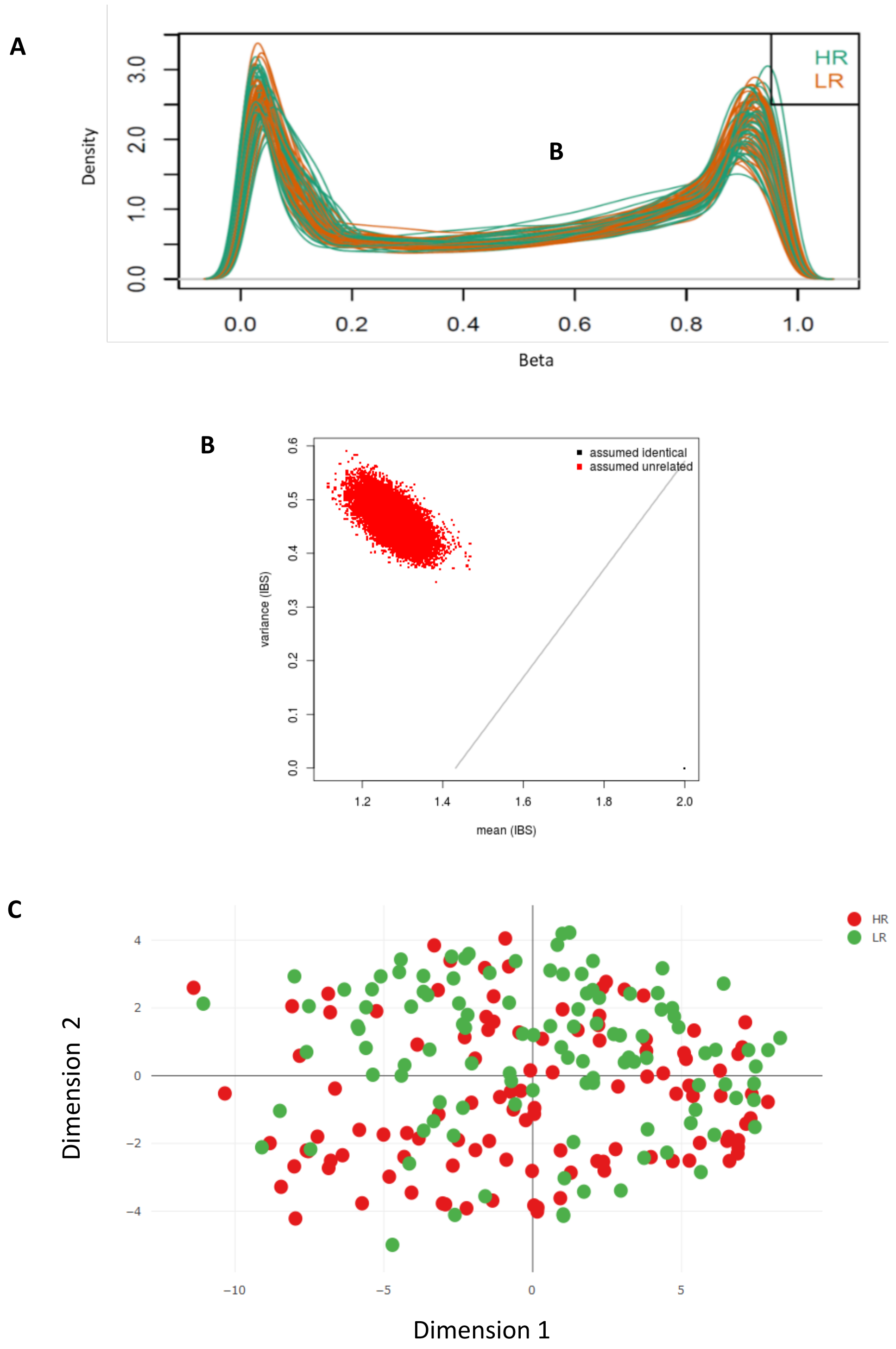


Abbreviations: high-risk bowel adenomas (HR), identical by state (IBS), and low-risk bowel adenomas (LR)

**Figure S2**

**A**

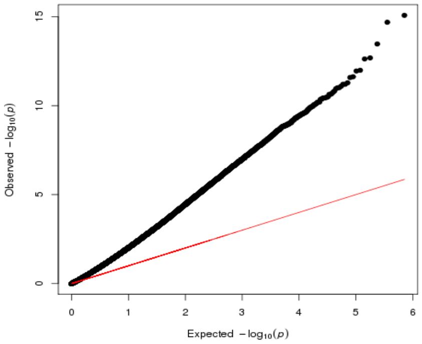


Lambda 2.10

**B**

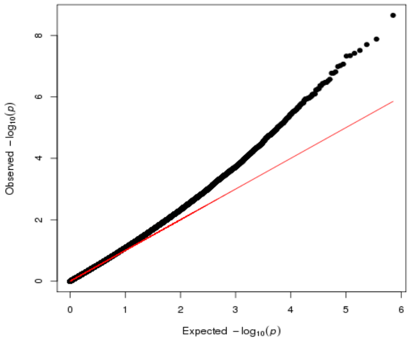


Lambda 1.09

**C**

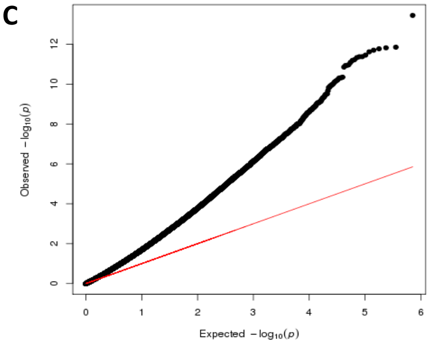


Lambda 1.89

**D**

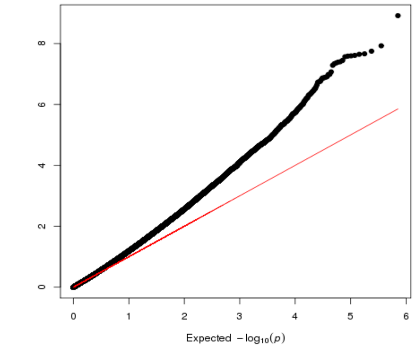


Lambda 1.09

**E**

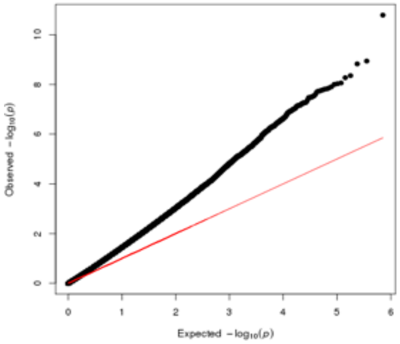


Lambda 1.62

**F**

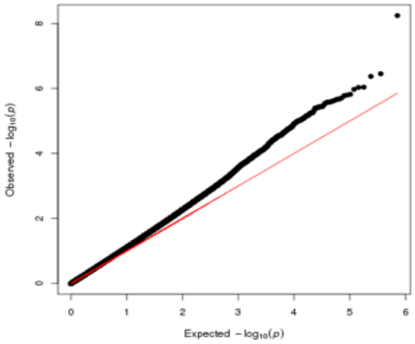


Lambda 1.15

Abbreviations: log (logarithm) and p-value (p)

### Figure S3

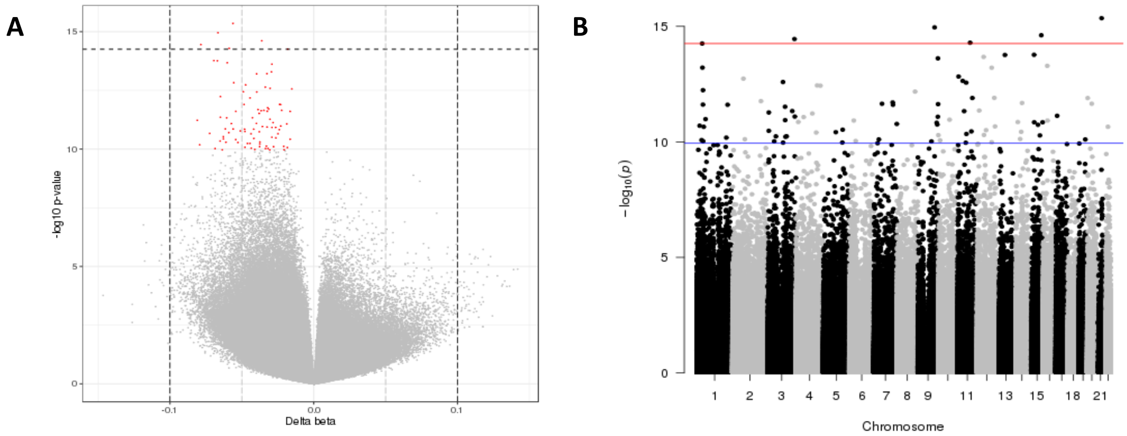


### Table S1

| Category | Details | n | (%) |
|---|---|---|---|
| Distance to concurrent CRC (mm) [1] | < 30 | 44 | (42) |
| | ≥ 30 | 39 | (37) |
| | ≥ 50 | 11 | (10) |
| | ≥ 100 | 12 | (11) |
| Number concurrent CRCs | 1 | 91 | (86) |
| | 2 | 15 | (14) |
| Concurrent CRC MS status | MSS | 25 | (24) |
| | Not tested | 81 | (76) |
| Number adenomas [2] | 1 to 3 | 66 | (62) |
| | 4 to 10 | 39 | (37) |
| | Unknown | 1 | (1) |

Abbreviations: colorectal cancer (CRC), greater than or equal to (≥), less than (<), millimetre (mm), microsatellite (MS), microsatellite stable (MSS), number (n) and percentage (%)

[1] Distance from case high-risk adenoma to closest concurrent CRC if more than one

[2] Case high-risk adenoma plus concurrent adenomas in each patients' high-risk bowel

**Table S2**

| | | DMPs | |
|---|---|---|---|
| Category | Gene | Putative function [1] | Cancer and lifestyle association details |
| CRC and/or adenomas | ANK2 (hyper, body) | Encodes a protein involved in the structural constituent of cytoskeleton, and enzyme binding. | Sometimes genetically altered in colorectal lateral spreading tumours (76) and gene mRNA may be core genes in **CRC** and potentially **progression** (77). |
| | ANXA6 (hypo, TSS200) | Encodes a protein involved with calcium ion and GTP binding, and transport of cholesterol. | Protein expression in stools may be biomarker for high-risk adenomas (65). Promoter hypermethylated and reduced expression in gastric cancer cell lines versus normal (78) and associated with obesity (79). |
| | CEMIP (hyper, body) | Encodes a protein involved in enzyme activity, metabolism, regulating **epithelial-mesenchymal transition**, tumour cell growth, invasion and **cancer** dissemination. | Hypomethylated in **adenomas** versus normal bowel tissue (64), and associated with the Wnt signalling pathway (80). Higher expression in **CRC** is associated with **poorer survival** (80). |
| | GUCY2C (hyper, body) | Encodes a protein involved in signal transduction, enzyme activity, **digestion,** and metabolism and uptake and actions of bacterial toxins pathways. | Postulated related to obesity association with **cancer risk** (81) with high caloric induced obesity associated with silencing of GUCY2C signalling in **colon epithelial** cells and thought to lead to **CRC** (63,64). Hypermethylation in ovarian cancer associated with longer survival (82). |
| | TPP1 (hyper, 3'UTR) | Encodes a protein involved in enzyme activity, lipid metabolic process and associated with **telomere length.** | Genomic alteration associated with telomere dysfunction increases **risk** of **CRC** (66). |
| Other cancers | DIAPH3 (hyper, body) | Encodes a protein involved in actin remodelling, enzyme activity and **ERK signalling**. | Increased expression in hepatocellular cancer, associated with **beta-catenin/TCF** signalling pathway and **epithelial-mesenchymal transition** (51). SNPs associated with **smoking measurement** (ever regular versus never regular) (83). |
| | RGS3 (hyper, 5'UTR) | Encodes a protein involved in enzyme activity, **MAPK** and **Wnt/β-catenin** signalling and **epithelial-mesenchymal transition**. | Increased expression associated with gastric cancer versus normal, poorer survival and aggressive cancer outcomes (52). |
| | TMEM135 (hyper, body) | Encodes a protein involved in **response to food**, and mitochondrial metabolism. | Near SNP associated with increased contralateral breast cancer risk (53) |

Abbreviations: ankyrin 2 (ANK2), annexin A6 (ANXA6), cell migration-inducing and hyaluronan-binding protein (CEMIP), colorectal cancer (CRC), diaphanous related formin 3 (DIAPH3), differentially methylated position (DMP), extracellular signal-regulated kinase (ERK), guanosine triphosphate (GTP), guanylate cyclase 2C (GUCY2C), hypermethylated (hyper), hypomethylated (hypo), messenger ribonucleic acid (mRNA), mitogen-activated protein kinase (MAPK), regulator of G protein signalling 3 (RGS3), single nucleotide polymorphism (SNP), T-cell factor (TCF), transcription start site (TSS), transmembrane protein 135 (TMEM135), tripeptidyl peptidase 1 (TPP1), untranslated region (UTR) and wingless-type (Wnt)

**Table S3**

| Details | | DMRs (n = 217) | | | | | | SM DMRs (n = 138) | | | | JR DMRs (n = 79) | | | |
|---|---|---|---|---|---|---|---|---|---|---|---|---|---|---|---|
| Chr | Gene | CpGs (n) | Length (bp) | Min FDR [1] | Stouffer p-value | Max db [2] | Mean db [3] | Length (bp) | Min FDR [1] | Stouffer p-value | Mean db [3] | Length (bp) | Min FDR [1] | Stouffer p-value | Mean db [3] |
| **1** | **GPX7** | **13** | **1469** | **2.27e-23** | **5.89e-11** | **10.5** | **5.2** | **1469** | **1.87e-11** | **3.01e-03** | **4.8** | **303** | **1.15e-07** | **9.41e-02** | **8.4** |
| 5 | MCIDAS | 7 | 669 | 1.65e-11 | 6.84e-07 | 11.2 | 8.5 | / | / | / | / | 669 | 1.54e-11 | 9.65e-04 | 15.4 |
| 8 | XKR6 | 15 | 1270 | 4.94e-12 | 6.85e-06 | 9.3 | 6.5 | / | / | / | / | 1270 | 8.00e-18 | 1.75e-04 | 12.0 |
| 5 | CCL28 | 11 | 815 | 2.95e-10 | 1.29e-05 | 10.6 | 7.0 | / | / | / | / | / | / | / | / |
| 22 | CECR2 | 8 | 1063 | 2.17e-10 | 7.29e-05 | -11.8 | -5.6 | / | / | / | / | / | / | / | / |
| 10 | FAM24B | 14 | 1582 | 1.16e-09 | 9.24e-05 | -12.1 | -7.4 | / | / | / | / | / | / | / | / |
| 2 | LBX2 | 11 | 843 | 1.97e-09 | 1.02e-04 | 10.1 | 6.5 | 688 | 3.42e-04 | 1.20e-01 | 6.0 | / | / | / | / |
| 11 | LUZP2 | 12 | 577 | 1.17e-08 | 1.24e-04 | 8.3 | 5.8 | / | / | / | / | / | / | / | / |
| 3 | VILL | 7 | 597 | 3.28e-06 | 1.10e-03 | -10.5 | -6.7 | 597 | 1.54e-06 | 3.75e-03 | -8.4 | / | / | / | / |
| 16 | SHISA9 | 7 | 1589 | 8.20e-07 | 3.41e-03 | -10.0 | -5.0 | / | / | / | / | 1421 | 1.03E-09 | 6.79e-03 | -9.6 |

Abbreviations: base-pair (bp), cat eye syndrome chromosome region candidate 2 (CECR2), C-C motif chemokine ligand 28 (CCL28), chromosome (chr), cytosine-phosphate-guanine (CpG), delta beta percentage (db), differentially methylated region (DMR), exponential notation (e), false discovery rate q-values (FDR), family with sequence similarity 24 member B (FAM24B), glutathione peroxidase 7 (GPX7), John Radcliffe Hospital (JR), ladybird homeobox 2 (LBX2), leucine zipper protein 2 (LUZP2), maximum (max), minimum (min), multiciliate differentiation and DNA synthesis-associated cell cycle (MCIDAS), number (n), shisa family member 9 (SHISA9), St Mark's Hospital (SM), villin like (VILL) and XK related 6 (XKR6)

[1] Minimum FDR of the smoothed estimate

[2] Maximum delta beta is the maximum delta beta within DMR

[3] Mean delta beta is the mean delta beta across DMR

**Table S4**

| | | DMRs | |
|---|---|---|---|
| Category | Gene | Putative function [1] | Cancer and lifestyle association details |
| CRC and/or adenomas | CCL28 (hyper) | Encodes a protein involved in chemokine and cytokine activity, immune response, signal transduction and **TCFß** signalling pathway. Expressed in **epithelium colon** and other **epithelium.** | Reduced expression in left **colon** cancer versus normal bowel tissue (68), and over-expression associated with breast **cancer progression** variables (84). |
| | MCIDAS (hyper) | Encodes a protein involved in protein binding, transcription coactivator activity, regulation of cell cycle and generation of multiciliated cells in respiratory **epithelium.** | Hypermethylated in serrated cancer versus normal bowel tissue (67). |
| | PITX2 (hyper) | Encodes a protein involved in transcription factor activity and the TGFB signalling pathway. | Promoter hypermethylation in CRC versus normal bowel tissue and lower CRC expression associated with **poorer survival**, however opposite for all these findings in other study (69,70). Promoter hypermethylation in prostate cancer correlated with increased cancer outcomes and predictor biochemical recurrence (85). |
| | UCP1 (hyper) | Encodes a protein involved in mitochondrial and ion transport, thermogenic respiration, temperature, **diet variations** and **regulation of energy balance.** Expressed in brown **adipose tissue.** | Positive expression in **CRC** versus normal tissue, and increased expression in CRC associated with better survival (86). |
| Other cancers | GPX7 (hyper) | Encodes a protein that protects against oxidative **DNA damage, double-strand breaks** and oesophageal **epithelia** from damage. | More promoter hypermethylation and reduced expression in gastric cancer versus normal gastric tissue (54). |

Abbreviations: C-C motif chemokine ligand 28 (CCL28), colorectal cancer (CRC), deoxyribonucleic acid (DNA), differentially methylated region (DMR), glutathione peroxidase 7 (GPX7), hypermethylation (hyper), multiciliate differentiation and DNA synthesis-associated cell cycle (MCIDAS), paired like homeodomain 2 (PITX2), transforming growth factor beta (TGFß) and uncoupling protein 1 (UCP1)

**Table S5**

**A**

| GO number | Ont | Term | N | DE | P.DE | BH FDR [1] |
|---|---|---|---|---|---|---|
| GO:0044424 | intracellular part | CC | 13186 | 9923 | 1.62e-98 | 3.58e-94 |
| GO:0005622 | intracellular | CC | 13457 | 10098 | 8.55e-96 | 9.45e-92 |
| GO:0005515 | protein binding | MF | 10667 | 8150 | 1.55e-87 | 1.15e-83 |
| GO:0043226 | organelle | CC | 12452 | 9347 | 7.94e-87 | 4.39e-83 |
| GO:0043229 | intracellular organelle | CC | 11552 | 8721 | 2.52e-85 | 1.11e-81 |
| GO:0043227 | membrane-bounded organelle | CC | 11540 | 8694 | 5.94e-84 | 2.19e-80 |
| GO:0005737 | cytoplasm | CC | 10613 | 8063 | 1.61e-80 | 5.09e-77 |
| GO:0043231 | intracellular membrane organelle | CC | 9926 | 7560 | 1.98e-80 | 5.48e-77 |
| GO:0005488 | binding | MF | 13685 | 10125 | 1.03e-75 | 2.54e-72 |
| GO:0003674 | molecular function | MF | 15933 | 11531 | 4.14e-71 | 9.17e-68 |
| GO:0044422 | organelle part | CC | 8575 | 6572 | 2.99e-70 | 6.01e-67 |
| GO:0044446 | intracellular organelle part | CC | 8372 | 6423 | 2.08e-69 | 3.83e-66 |
| GO:0005623 | cell | CC | 15295 | 11130 | 3.18e-68 | 5.41e-65 |
| GO:0044464 | cell part | CC | 15265 | 11110 | 1.21e-67 | 1.92e-64 |
| GO:0044444 | cytoplasmic part | CC | 8851 | 6739 | 4.62e-65 | 6.82e-62 |
| GO:0044237 | cellular metabolic process | BP | 10027 | 7501 | 8.02e-65 | 1.11e-61 |
| GO:0008152 | metabolic process | BP | 10714 | 7968 | 1.22e-64 | 1.58e-61 |
| GO:0044238 | primary metabolic process | BP | 9911 | 7377 | 6.22e-59 | 7.65e-56 |
| GO:0071704 | organic substance metabolic process | BP | 10264 | 7625 | 9.26e-59 | 1.08e-55 |
| GO:0008150 | biological process | BP | 15955 | 11456 | 2.05e-58 | 2.26e-55 |

**B**

| Pathway | N | DE | P.DE | BH FDR [1] |
|---|---|---|---|---|
| Metabolic pathways | 1410 | 1061 | 6.01e-09 | 2.03e-06 |
| Axon guidance | 174 | 154 | 1.99e-06 | 3.36e-04 |
| Ubiquitin mediated proteolysis | 132 | 115 | 7.15e-06 | 6.55e-04 |
| Autophagy - animal | 130 | 115 | 8.20e-06 | 6.55e-04 |
| Pathways in cancer | 506 | 401 | 9.69e-06 | 6.55e-04 |
| Signalling pathways regulating pluripotency of stem cells | 139 | 120 | 1.69e-05 | 9.54e-04 |
| ErbB signalling pathway | 80 | 74 | 2.35e-05 | 1.13e-03 |
| Bacterial invasion of epithelial cells | 76 | 70 | 3.47e-05 | 1.47e-03 |
| Rap1 signalling pathway | 207 | 175 | 5.25e-05 | 1.97e-03 |
| Endocrine resistance | 95 | 85 | 6.43e-05 | 2.17e-03 |
| Cell cycle | 121 | 104 | 7.07e-05 | 2.17e-03 |
| Thyroid hormone signalling pathway | 117 | 103 | 8.76e-05 | 2.47e-03 |
| RNA degradation | 76 | 67 | 1.12e-04 | 2.92e-03 |
| Alzheimer disease | 345 | 269 | 1.87e-04 | 4.23e-03 |
| Pathways of neurodegeneration - multiple diseases | 448 | 346 | 1.88e-04 | 4.23e-03 |
| Human papillomavirus infection | 319 | 255 | 2.14e-04 | 4.52e-03 |
| Focal adhesion | 193 | 163 | 2.51e-04 | 4.63e-03 |
| Endocytosis | 242 | 197 | 2.61e-04 | 4.63e-03 |
| Bladder cancer | 40 | 38 | 2.66e-04 | 4.63e-03 |
| cGMP-PKG signalling pathway | 161 | 135 | 2.74e-04 | 4.63e-03 |

**C**

| GO number | Ont | Term | N | DE | P.DE | BH FDR [1] |
|---|---|---|---|---|---|---|
| GO:0045120 | pronucleus | CC | 15 | 3 | 7.78e-05 | 1.00e+00 |
| GO:0005540 | hyaluronic acid binding | MF | 21 | 3 | 2.51e-04 | 1.00e+00 |
| GO:0046976 | histone methyltransferase activity | MF | 5 | 2 | 5.98e-04 | 1.00e+00 |
| GO:1900019 | regulation of protein kinase C activity | BP | 5 | 2 | 9.64e-04 | 1.00e+00 |
| GO:1900020 | positive regulation protein kinase C | BP | 5 | 2 | 9.64e-04 | 1.00e+00 |
| GO:0070317 | negative regulation G0 to G1 transition | BP | 34 | 3 | 1.30e-03 | 1.00e+00 |
| GO:0070316 | regulation of G0 to G1 transition | BP | 38 | 3 | 1.47e-03 | 1.00e+00 |
| GO:0045023 | G0 to G1 transition | BP | 40 | 3 | 1.56e-03 | 1.00e+00 |
| GO:0015679 | plasma membrane copper ion transport | BP | 1 | 1 | 1.72e-03 | 1.00e+00 |
| GO:0098705 | copper ion across plasma membrane | BP | 1 | 1 | 1.72e-03 | 1.00e+00 |
| GO:0097539 | ciliary transition fiber | CC | 10 | 2 | 2.87e-03 | 1.00e+00 |
| GO:0099515 | actin filament-based transport | BP | 9 | 2 | 2.97e-03 | 1.00e+00 |
| GO:0044446 | intracellular organelle part | CC | 8372 | 61 | 3.40e-03 | 1.00e+00 |
| GO:0044422 | organelle part | CC | 8575 | 62 | 3.79e-03 | 1.00e+00 |
| GO:0031406 | carboxylic acid binding | MF | 176 | 5 | 3.80e-03 | 1.00e+00 |
| GO:0043177 | organic acid binding | MF | 177 | 5 | 3.84e-03 | 1.00e+00 |
| GO:0035518 | histone H2A monoubiquitination | BP | 15 | 2 | 3.87e-03 | 1.00e+00 |
| GO:0008465 | glycerate dehydrogenase activity | MF | 1 | 1 | 4.03e-03 | 1.00e+00 |
| GO:0042470 | melanosome | CC | 104 | 4 | 4.24e-03 | 1.00e+00 |
| GO:0048770 | pigment granule | CC | 104 | 4 | 4.24e-03 | 1.00e+00 |

**D**

| Pathway | N | DE | P.DE | BH FDR [1] |
|---|---|---|---|---|
| Mineral absorption | 54 | 2 | 3.12E-02 | 1.00e+00 |
| MicroRNAs in cancer | 297 | 4 | 5.19E-02 | 1.00e+00 |
| Lysine degradation | 61 | 2 | 6.47E-02 | 1.00e+00 |
| Renal cell carcinoma | 65 | 2 | 8.17E-02 | 1.00e+00 |
| Linoleic acid metabolism | 29 | 1 | 9.09E-02 | 1.00e+00 |
| Selenocompound metabolism | 17 | 1 | 9.19E-02 | 1.00e+00 |
| alpha-Linolenic acid metabolism | 25 | 1 | 1.10E-01 | 1.00e+00 |
| Glycerophospholipid metabolism | 93 | 2 | 1.24E-01 | 1.00e+00 |
| Glycine, serine and threonine metabolism | 35 | 1 | 1.29E-01 | 1.00e+00 |
| RNA polymerase | 29 | 1 | 1.32E-01 | 1.00e+00 |
| Glycosaminoglycan biosynthesis - heparan sulfate/heparin | 23 | 1 | 1.41E-01 | 1.00e+00 |
| Glyoxylate and dicarboxylate metabolism | 30 | 1 | 1.43E-01 | 1.00e+00 |
| SNARE interactions in vesicular transport | 32 | 1 | 1.50E-01 | 1.00e+00 |
| Lysosome | 123 | 2 | 1.63E-01 | 1.00e+00 |
| Cytosolic DNA-sensing pathway | 55 | 1 | 1.72E-01 | 1.00e+00 |
| Pyruvate metabolism | 38 | 1 | 1.83E-01 | 1.00e+00 |
| Regulation of actin cytoskeleton | 207 | 3 | 1.91E-01 | 1.00e+00 |
| Arachidonic acid metabolism | 63 | 1 | 1.94E-01 | 1.00e+00 |
| Retinol metabolism | 62 | 1 | 1.94E-01 | 1.00e+00 |
| Neurotrophin signalling pathway | 114 | 2 | 2.02E-01 | 1.00e+00 |

Abbreviations: benjamini-hochberg false discovery rate q-values (BH FDR), biological process (BP), cellular component (CC), cyclic guanosine monophosphate-protein kinase G (cGMP-PKG), deoxyribonucleic acid (DNA), epidermal growth factor receptor (ErbB), exponential notation (e), gene ontology (GO), molecular function (MF), number of genes differentially methylated (DE), number of genes in the term (N), ontology (ont), p-value for over-representation of the term (P.DE), ras-related protein 1 (Rap1), ribonucleic acid (RNA) and soluble N-ethylmale-imide-sensitive factor-attachment protein receptor (SNARE)

### Supplementary Figure and Table legends

**Figure S1 Sample data quality control and clustering plots.** The following quality control plots were created for the high-risk (HR) and low-risk (LR) bowel adenoma EPIC methylation BeadChip data **a)** Beta density plot representative of the John Radcliffe Hospital samples beta distributions after probe filtering and beta-mixture quantile normalisation (BMIQ) **b)** Sample relationships plot calculated from 720 single nucleotide polymorphisms (SNP) from the samples EPIC methylation BeadChips **c)** Multidimensional scaling plot of the BMIQ normalised and batch effect corrected sample beta values clustered by 1000 most variable methylated cytosine-phosphate-guanine (CpG) sites.

**Figure S2 Differentially methylated positions plots.** The following quantile-quantile (Q-Q) plots were created for the high-risk (HR) compared to low-risk (LR) bowel adenoma differentially methylated position (DMPs) adjusted for confounders. Where, the black dots are the observed p-values, and the red lines are the expected p-value distributions. The lambdas represent the estimated genomic inflation calculated from the DMP p-values. **a)** Q-Q plot using DMP p-values **b)** Q-Q plot using DMP p-values after genomic inflation correction **c)** Q-Q plot using stratified St Mark's Hospital DMP p-values **d)** Q-Q plot using stratified St Mark's Hospital DMP p-values after genomic inflation correction **e)** Q-Q plot using stratified John Radcliffe Hospital DMP p-values **f)** Q-Q plot using stratified John Radcliffe Hospital DMP p-values after genomic inflation correction

**Figure S3 Differentially methylated positions volcano and manhattan plots**
The following plots were created for the high-risk (HR) compared to low-risk (LR) bowel adenoma differentially methylated positions (DMPs) adjusted for confounders and genomic inflation corrected **a)** Volcano plot, where the x-axis represents the DMP delta beta values, and the y-axis represents the DMP p-values. The left and right side of the x-axis represents hypomethylation and hypermethylation of the DMPs, respectively. The vertical dashed black lines marks the DMPs with a delta beta ≤ - 0.10 or ≥ 0.10. DMPs coloured in red have Benjamini-Hochberg false discovery rate (BH FDR) q-values < 0.05, and the horizontal dashed black line marks the Bonferroni p-value significance threshold. **b)** Manhattan plot, where the x-axis represents the chromosomes in the human genome and the y-axis represents the DMP p-values. DMRs above the blue horizontal line have BH FDR q-values < 0.05, and the horizontal red line marks the Bonferroni genome-wide p-value significance threshold.

**Table S1 Patients high-risk bowel characteristics.** The patients' high-risk (HR) bowel characteristics are shown in more detail.

**Table S2 Differentially methylated position genes and literature.** For DMPs from Table 2, selected gene functions and then lifestyle exposures and cancer associations are shown. DMP genes with Bonferroni significant p-values $< 7 \times 10^{-8}$ are highlighted in red. Gene functions and associations were sourced from the GeneCards Human Gene database and the literature. Key associations are highlighted in bold.

**Table S3 Differentially methylated regions with principal component adjustment.** High-risk (HR) compared to low-risk (LR) bowel adenoma differentially methylated regions (DMRs) adjusted for confounders and principal components 1 and 2, with mean delta beta values ≥ 5%, Stouffer p-values < 0.05 and a minimum of 7 cytosine-phosphate-guanines (CpGs) are shown. These were ordered by decreasing Stouffer p-values, and then matched to their corresponding stratified St Mark's Hospital and John Radcliffe Hospital study-site DMR values. The significant replicated study-site DMR is highlighted in bold.

**Table S4 Differentially methylated region genes and literature.** For DMRs from Tables 3 and S3, selected gene functions and then lifestyle exposures and cancer associations are shown. Gene functions and associations were sourced from the GeneCards Human Gene database and the literature. Key associations are highlighted in bold.

**Tables S5 Gene ontology and pathway enrichment analysis.** The following enrichment analyses were carried out for the high-risk (HR) compared to low-risk (LR) bowel adenoma differentially methylated positions (DMPs) that were adjusted for confounders and genomic inflation corrected, with p-values > 0.05 **a)** Gene Ontology (GO) analysis and ordered by decreasing enrichment Benjamini-Hochberg false discovery rate (BH FDR) q-values **b)** Kyoto Encyclopaedia of Genes and Genomes (KEGG) pathway analysis and ordered by decreasing enrichment BH FDR q-values. The following enrichment analyses were carried for these HR compared to LR bowel adenoma DMPs, with BH FDR q-values > 0.05 **c)** GO analysis and ordered by decreasing enrichment BH FDR q-values **d)** KEGG pathway analysis and ordered by decreasing enrichment BH FDR q-values.

### Supplementary datasets and further details

#### High and low-risk bowel adenoma epigenome-wide association study

*Study design*

Patients were classified as having an inherited CRC syndrome if this was listed in their pathology records or if their pathology report histories met the definitions of the inherited CRC syndromes (4,5,87). Only patients HR bowel adenomas with no more than 10 adenomas within 6 months either side of the study adenoma and patients LR bowel adenomas with no more than 10 adenomas in the pathology records history were included in this study, to help reduce the likelihood that some of the study HR and LR bowel adenomas could have undiagnosed inherited polyposis. For each patient, clinical and sample data was collated from their hospital pathology records. Where patients gender was defined as male or female, age in 5-year groupings at adenoma diagnosis and adenoma bowel-side as left or right, lesion-size in mm's, dysplasia as LGD or HGD, and architecture as TA, TVA, or VA.

*Laboratory*

From the patients' adenoma samples, at least 650ng DNA was extracted using the using the Covaris truXTRAC FFPE DNA extraction kit. Then checked for DNA quality using a Qubit 3.0 fluorometer and Illumina FFPE QC by quantitative polymerase chain reaction (qPCR) using the FFPE QC qPCR assay kit. Samples were then normalised and ligated using the REPLI-g FFPE kit based on the Thirlwell methodology (74), and bisulfite converted using the Zymo EZ DNA Methylation Kit, with an extended polymerase chain reaction (PCR) cycle for increased bisulfite conversion efficiency. The DNA bisulfite conversion efficiency was tested by using a qPCR assay. Samples that had a DNA concentration ≥ 650 ng and a FFPE qPCR Cq value ≥ 6 were deemed suitable for DNA methylation EPIC BeadChip profiling.

#### External high and low-risk normal bowel tissue epigenome-wide association studies

The open-source Gene Expression Omnibus (GEO) GSE132804 HR and LR normal bowel tissue datasets (31) were downloaded and used for validation. This consisted of the GSE132804 - GPL21145 EPIC and the GSE132804 - GPL13534 450K methylation BeadChip EWAS datasets. For both datasets, the HR normal bowel samples were collected by surgical resection from newly diagnosed CRC patients in the ColoCare study (88) (age 19–85), and the LR normal bowel samples by endoscopic biopsy from patients undergoing colonoscopies (age 19–85) in the GICaRes study

(89). Following the protocols approved by the Institutional Review Board for the University of Washington Medical Center (Seattle, United States of America).
